# The emergence, maintenance and demise of diversity in a spatially variable antibiotic regime

**DOI:** 10.1101/158337

**Authors:** Alanna M. Leale, Rees Kassen

**Author notes:** Corresponding author: Alanna M. Leale.

## Abstract

Antimicrobial resistance (AMR) is a growing global threat that, in the absence of new antibiotics, requires effective management of existing drugs. Here, we explore how changing patterns of drug delivery modulates the spread of resistance in a population. Resistance evolves readily under both temporal and spatial variation in drug delivery and fixes rapidly under temporal, but not spatial, variation. Resistant and sensitive genotypes coexist in spatially varying conditions due to a resistance-growth rate trade-off which, when coupled to dispersal, generates negative frequency-dependent selection and a quasi-protected polymorphism. Coexistence is ultimately lost, however, because resistant types with improved growth rates in the absence of drug spread through the population. These results suggest that spatially variable drug prescriptions can delay but not prevent the spread of resistance and provide a striking example of how the emergence and eventual demise of biodiversity is underpinned by evolving fitness trade-offs.

---

The effectiveness of antibiotic therapy to control infection is being steadily undermined by the combination of divestment in drug discovery, widespread occurrence of genetic resistance among microbes isolated from natural environments^1^, and continued evolution of resistant strains among all major pathogens^2^. Managing our existing arsenal of drugs to ensure they remain effective for as many people for as long as possible is therefore an urgent public health priority.

Beyond a blanket ban on prescriptions, there is little consensus on how to delay or prevent the emergence and spread of resistance. Using multiple drugs with distinct cellular and genetic targets is often suggested as the most effective treatment but sequential use can often select for multi-drug resistant strains^3^, while simultaneous delivery (as a drug cocktail) can be difficult for patients to tolerate^4^. An alternative approach is to exploit the use of drug-free sanctuaries or refuges in time or space to make it more difficult for selection to fix resistant strains. Sensitive strains that do not pay a fitness cost of resistance^5^ are expected to outcompete resistant strains in drug sanctuaries, implying that resistance could be kept at manageably low levels if dispersal introduces sensitive strains at a sufficiently high rate^6–8^. Sanctuaries have been used to good effect in managing resistance in agricultural systems^9,10^but their role in governing the spread of resistance in health settings remains unclear^11,12^.

Theory suggests that the way sanctuaries are experienced in time and space can profoundly impact resistance evolution^7,11,13,14^. Periodic delivery of a drug in time generates regular cycles of strong antibiotic selection followed by periods of relaxed selection when the drug is either metabolized, excreted, or not in use^15,16^. Intermittent dosing or ward-wide use generates fluctuating selection that leads to the evolution of broadly adapted resistant types with high fitness in both the presence and absence of drug^17^. Drug delivery may also be spatially variable across wards in a hospital or among compartments (tissues or organs) within a host, generating divergent selection that can lead to the emergence of a single resistant generalist or, if selection is sufficiently strong relative to dispersal (for example, through the movement of patients and staff between hospital wards), the coexistence of more narrowly-adapted niche specialists that trade-off drug resistance with growth rate in the absence of drug^7,11,13^.

To evaluate the impact of drug sanctuaries on the emergence and spread of AMR, we tracked the evolution of resistance and fitness in twelve independently evolved, isogenic lines of *P. aeruginosa* strain PA14 propagated in environments that varied through time or space with sub-inhibitory concentrations of the commonly used fluoroquinolone antibiotic, ciprofloxacin. *P. aeruginosa* ranks among the top three drug resistant pathogens globally^18^ and causes both acute infections of wounds and chronic infections of the respiratory tract, where it is a major source of morbidity and mortality in adults with cystic fibrosis (CF)^19,20^. We use drug concentrations that resemble that found in the sputum of CF patients undergoing fluoroquinolone treatment during exacerbation^21^. Controls were a permissive (PERM) environment consisting of drug free Luria-Bertrani (LB) medium and a constant selective environment (CONS) comprised of LB supplemented daily with sub-inhibitory concentrations (0.3 ug/ml) of ciprofloxacin sufficient to reduce population densities to 20% of that in the absence of drug. Temporally varying environments (TEMP) were constructed by transferring aliquots from each evolving population into either permissive (no drug) or selective (0.3ug/ml) conditions on alternating days. Spatially variable environments (SPAT) consisted of two subpopulations, or patches, one permissive and one selective, with equal volume aliquots from each mixed and redistributed into fresh drug-free and drug-supplemented media daily (see Methods and Supplementary Fig. S1). The SPAT treatment resembles a well-known model in population genetics in which genetic polymorphism can be maintained through negative frequency dependent selection, provided there is a strong fitness trade-off among genotypes across patches and population regulation occurs at the level of the local patch rather than the total population^11,22–24^. In contrast to previous work^25–28^, we do not explicitly manipulate either fitness trade-offs or the manner of population regulation but instead allow these variables to evolve naturally over the course of the experiment.

## Results and Discussion

After 20 days (∼133 generations) of selection resistance failed to evolve in the absence of drug selection (Fig. 1A) but did evolve under all other conditions (Fig. 1B), Fig. 1D). The level of resistance was assayed by determining the minimum inhibitory concentration (MIC; the lowest drug concentration that completely inhibits growth) for eight randomly selected colonies from each evolved population. Resistance fixed in all populations experiencing drug sanctuaries in time (Fig. 1C) but not space, where 6 of 10 populations contained a mixture of resistant and sensitive colonies (Fig. 1D; note that two populations in this treatment were lost due to contamination). Notably, the least resistant genotypes in the spatially structured environment (Fig. 1D) were as sensitive to ciprofloxacin as the ancestor and evolved populations from the permissive environment (Fig. 1A), while the most resistant genotypes had similar MICs as evolved genotypes from both the constant (Fig. 1B) and temporally varying (Fig. 1C) environments. Thus, drug sanctuaries do little to prevent the initial evolution of resistance, but when spatially structured, can slow or prevent the rate at which resistance sweeps through the population. More generally, divergent selection caused by spatial heterogeneity in antibiotic concentrations can promote diversification and coexistence between resistant and sensitive genotypes, while fluctuating selection on a similar time-scale does not.

**Figure 1.**
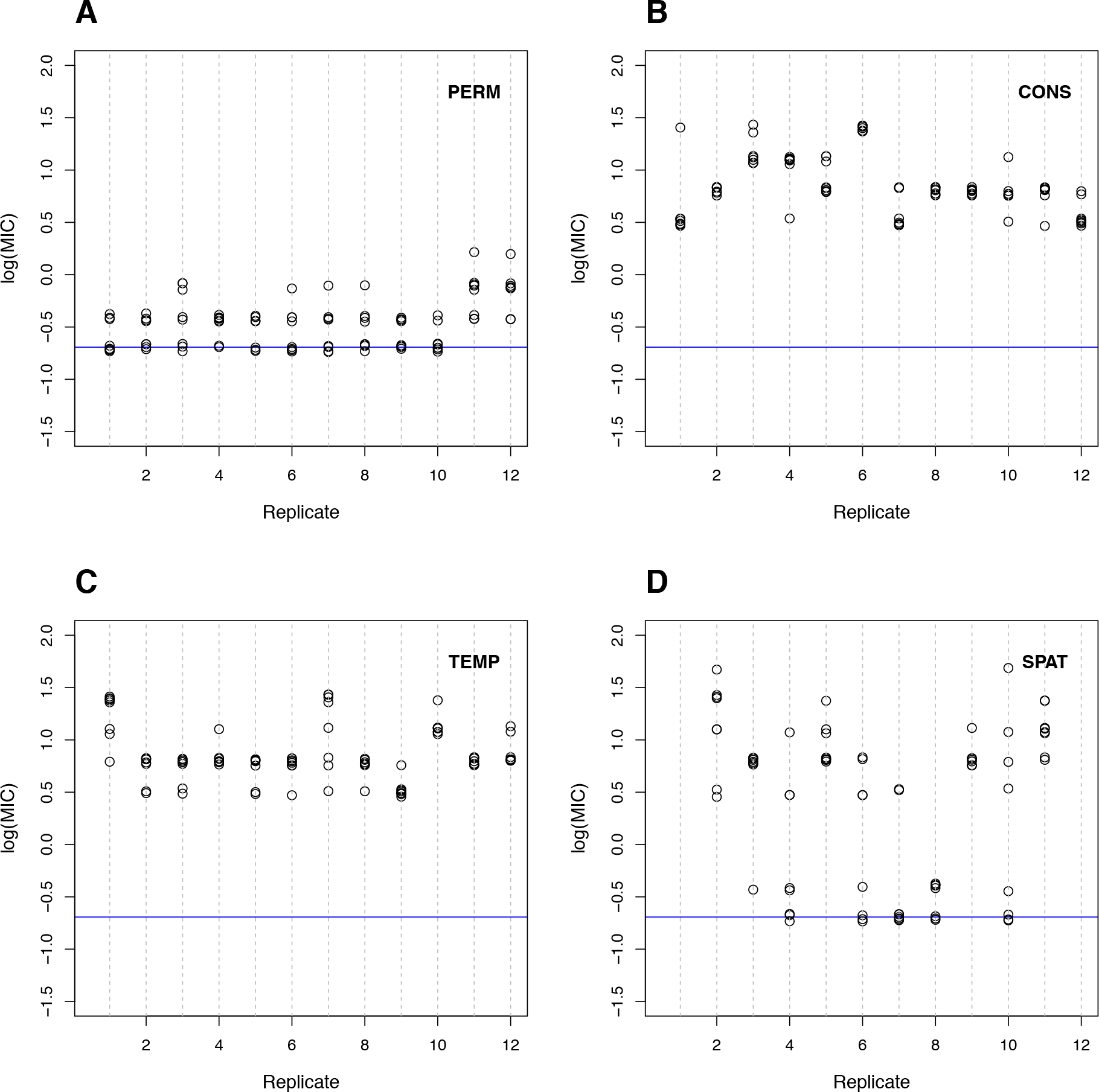
Resistance, measured as the log minimum inhibitory concentration (log(MIC)) of each of eight isolates from each evolved population after 20 days of serial transfer (∼133 generations). **(A)** PERM, **(B)** CONS, **(C)** TEMP, and **(D)** SPAT. Blue line indicates resistance level of ancestral PA14 isolate (MIC= 0.5ug/mL). An isolate is deemed resistant if its MIC > 2ug/mL (log(MIC) = 0.3). Data points are vertically jittered for clarity.

Does the diversity in resistance profiles in the spatially variable environment reflect stable coexistence, or might it simply be a transient effect reflecting a reduced overall strength of selection for resistant types^29^? Four lines of evidence point to diversity being stably maintained in the SPAT populations. First, coexistence between resistant and sensitive strains persists through an additional 20 days of selection in 8 of 10 populations from the SPAT treatment, albeit with marked fluctuations through time (Fig. 2). Second, we did not observe any evidence of sensitive genotypes persisting under temporally variable conditions, as might be expected if the primary effect of the permissive patch is to slow the rate of fixation of resistance mutations (Supplementary Fig. S2). Third, we observed the expected trade-off between resistance and growth rate in the absence of drug for the eight previously isolated colonies at day 20 and an additional 8 colonies isolated at day 40 (Fig. 3A, Supplementary Table S1; mixed linear analysis of covariance between growth in LB and log(MIC)*day, with population as a random effect: *slope = −0.3558, P < 0.0001*). Fourth, and most convincingly, invasion from rare experiments (see Methods) between four randomly-selected pairs of resistant and sensitive isolates from SPAT populations at day 20 and day 40 show that rare genotypes always have higher fitness than common ones, providing direct evidence for negative frequency dependent selection (Fig. 4, Supplementary Table S2; mixed linear analysis of covariance between relative fitness (ω) and day*resistance frequency, with random effects of individual isolate pair nested in population: *slope = −0.41871, P < 0.0001).* These results suggest that divergent selection leads to the stable coexistence of resistant and sensitive strains through negative frequency dependent selection, consistent with predictions from models for the maintenance of polymorphism in spatially variable environments^22–24^.

**Figure 2.**
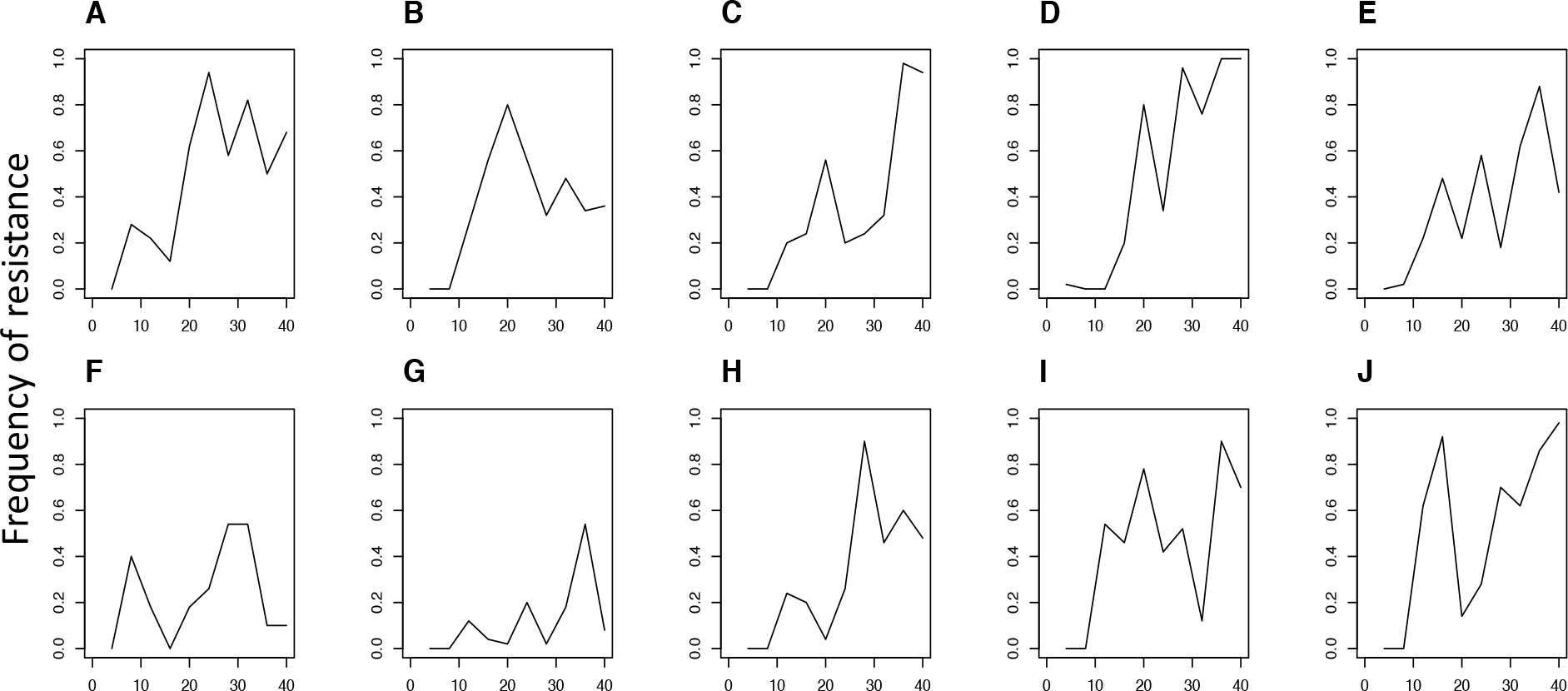
Dynamics of resistance within all SPAT populations. Each panel (A-J) is an independently evolved population. Sensitive colonies were detected at all time points in all populations except populations C, D, and J where all 50 colonies were resistant by day 36 and/or 40.

**Figure 3.**
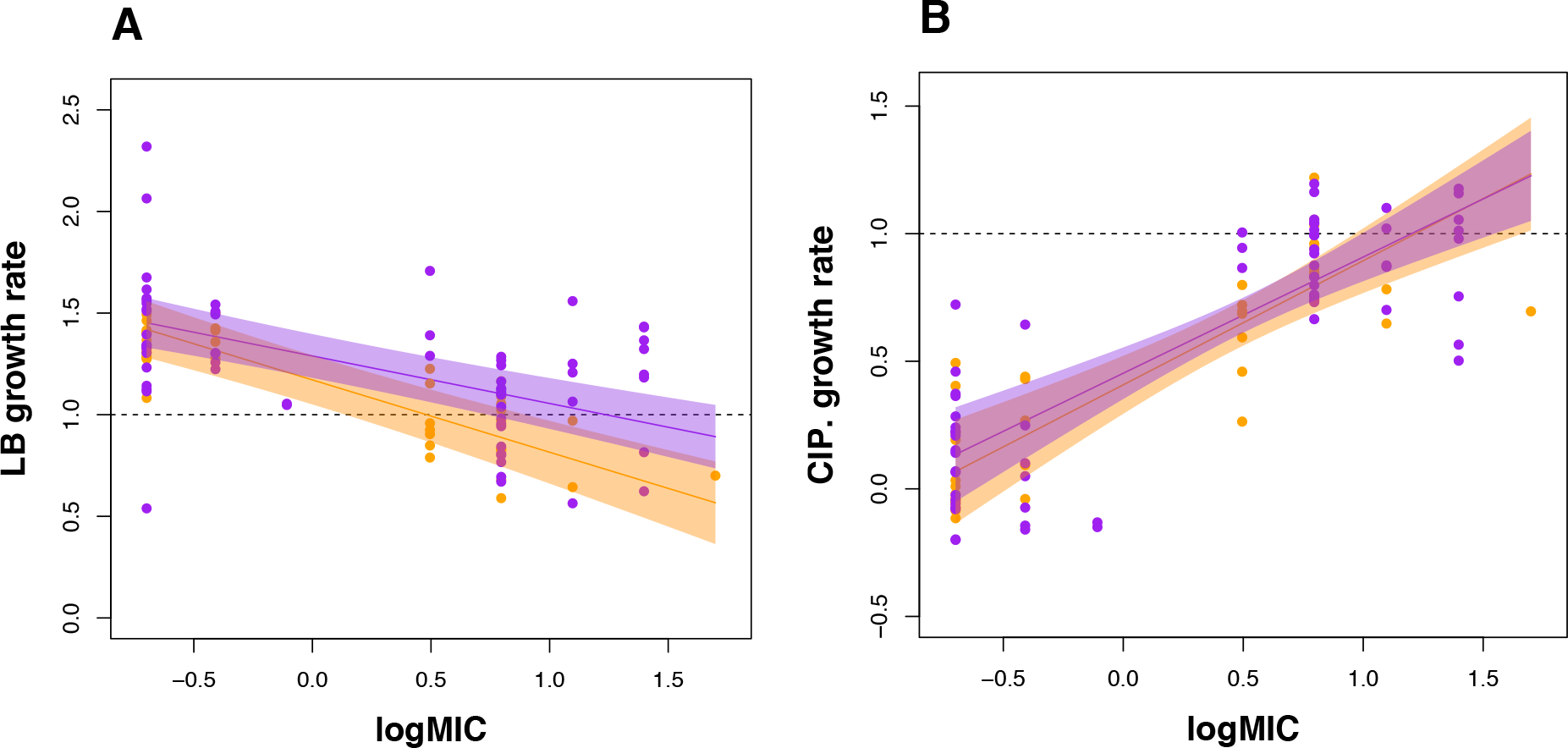
The trade-off between resistance and growth rate in LB (A) and LB supplemented with 0.3μg/mL ciprofloxacin (B). Data is standardized to the growth rate of the ancestral PA14 in LB (horizontal dashed lines in each panel). Shaded areas represent 95% confidence intervals for day 20 (orange) and day 40 (purple). Only populations with both resistant and sensitive isolated colonies were included for analyses.

**Figure 4.**
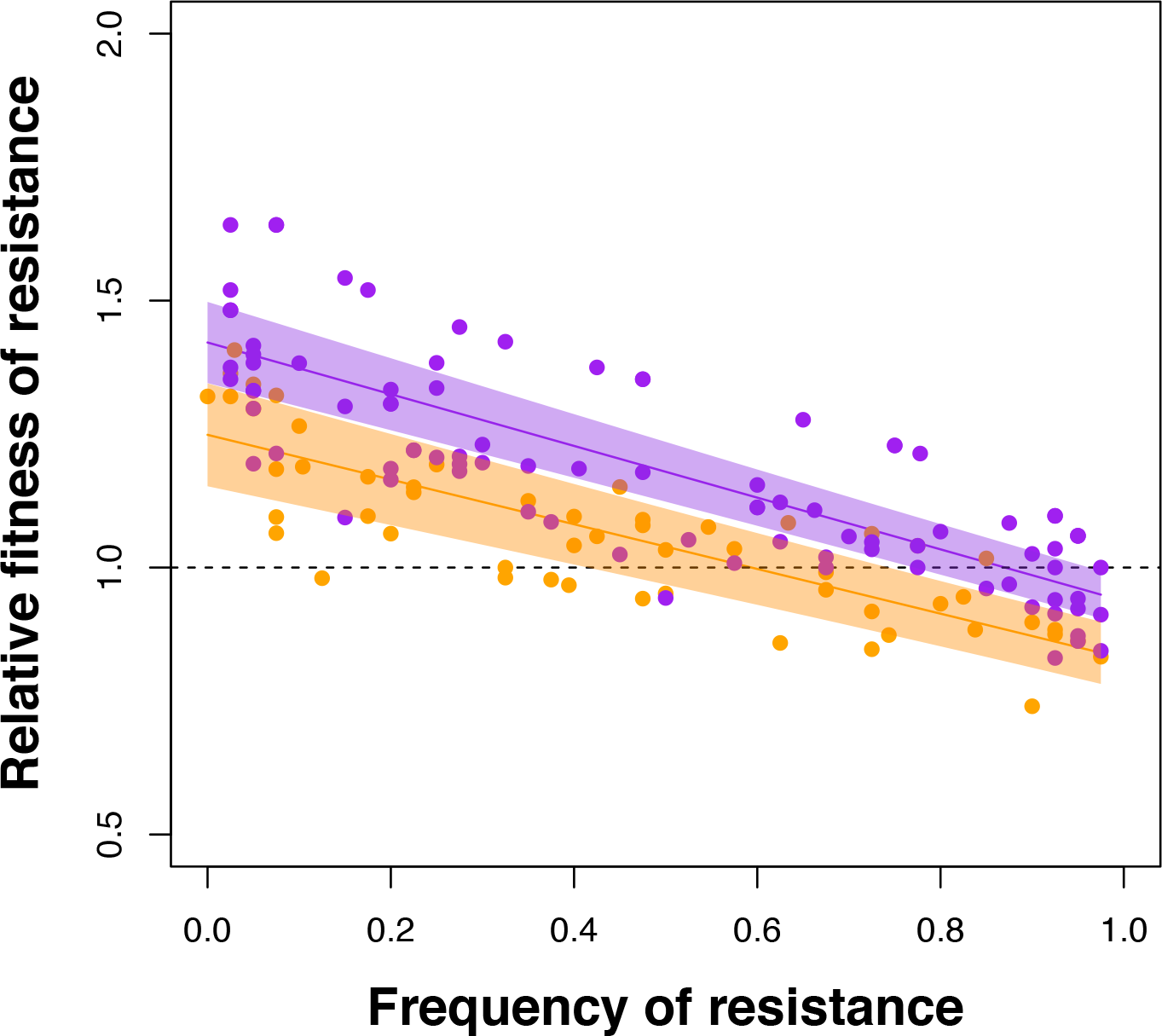
Negative frequency-dependent selection between resistant and sensitive strains within populations. Fitness of a resistant colony relative to a paired sensitive colony (३) is plotted as a function of its starting frequency. Shaded areas represent 95% confidence intervals for day 20 (orange) and day 40 (purple). The frequency of resistance at equilibrium is given by the intersection of the regression line with relative fitness (३) = 1. Only populations with both resistant and sensitive isolated colonies were included for analyses.

Despite the presence of strong negative frequency dependent selection, the polymorphism does not appear to be stable in our populations on evolutionary time scales. Coexistence persists at day 40 in just three populations (Fig. 2A, B, H) whereas four populations are fixed or nearly fixed for resistance (Fig. 2C, D, I, J) and nearly lost in two (Fig. 2F, G). The fixation or loss of resistant strains could be due to stochastic variation generated by time-lagged frequency dependent selection^30^, perhaps associated with the daily serial transfer protocol we imposed in our experiment, or from high mutation supply rates generating complex dynamics among competing strains (clonal interference). To distinguish between these alternatives, we tracked the frequency of resistance between three independent pairs of resistant and sensitive isolates over time under spatially variable conditions (see Methods). In the absence of clonal interference we expect the frequency of resistance to tend towards an internal equilibrium without fluctuations, regardless of starting frequency. As expected, the frequency of resistant genotypes converges towards an equilibrium point between 0 and 1 within the first two days and remains relatively stable until day 8, after which frequencies diverge again, presumably as mutations are reintroduced into the population (Fig.5). These results suggest that the fluctuations in the frequency of resistant and sensitive strains we observed in our original experiment are likely due to high mutation supply rates generating clonal interference (see also Maddemsetti *et al.* 2015^31^), rather than stochastic effects associated with our transfer protocol.

**Figure 5.**
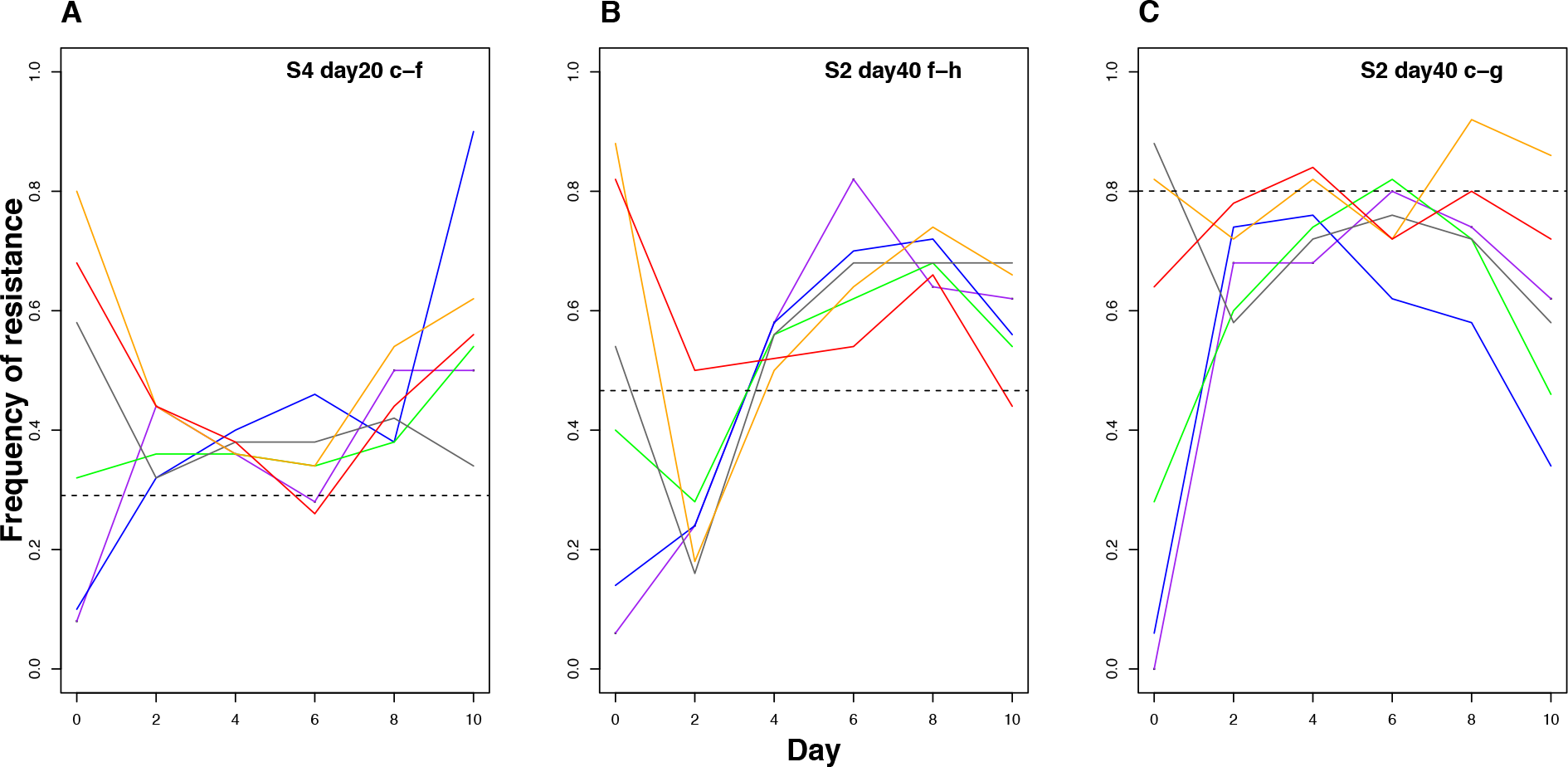
Dynamics of resistant-sensitive pairs starting from different starting frequencies. Panels A-C depict the relative frequency of distinct pairs resistant-sensitive strains over time for six independent populations begun from different starting frequencies (coloured lines). Dashed axes represent predicted stable equilibrium frequencies for each individual pair estimated from initial invasion from rare experiments reported in Supplementary Fig. S3.

High levels of clonal interference imply that there is a steady source of genetic variation on which selection can act, over and above the diversity supported by negative frequency dependent selection. If so, are these populations continuing to evolve or does negative frequency dependent selection act to prevent further evolution? To answer this question we focused attention on the how the trade-off between MIC and growth rate under drug-free conditions changed over the selection experiment. We find that the slope of this trade-off evolves to become shallower at day 40 than at day 20 (Supplementary Table S1; log(MIC)*day interaction: *F = 0.1221, P = 0.0382*). Inspection of Figure 3A suggests that this effect is due to increases in the growth rate of resistant isolates in LB, rather than loss of resistance (Fig. 3A). Further support for this interpretation comes from the lack of change in growth rate of resistant isolates in the presence of ciprofloxacin between days 20 and 40 (Fig. 3B, Supplementary Table S3; log(MIC)*day interaction: *F* = −0.0311, *P* = 0.6167). These results are consistent with the selection of second-site compensatory mutations that improve growth rate under drug-free conditions without compromising resistance^17,32–34^. As a consequence of this weakened trade-off, the internal equilibrium frequency of resistant strains (i.e., resistance frequency when ω = 1) increases on average from 59% at day 20 to 87% by day 40 (Fig. 4), implying that resistant genotypes are slowly spreading through most populations. A putatively stable polymorphism on ecological time scales can thus be readily undermined by selection which, in this case, leads to the evolution of broadly-adapted genotypes that are both resistant and have high fitness under drug-free conditions and the eventual loss of sensitive genotypes.

## Conclusion

Taken together, our results provide a rather bleak prognosis for the use of drug sanctuaries to manage antimicrobial resistance. Perhaps not surprisingly, given the strong selection generated by antibiotics, drug sanctuaries by themselves cannot prevent the evolution of drug resistance. Nevertheless, the manner in which drug sanctuaries are experienced by a pathogen can impact the rate at which resistance spreads in a population: while temporal variation in drug sanctuaries does little to prevent the rise of resistance, spatially distributed sanctuaries can slow the rate at which resistance fixes in a population. In line with the predictions of evolutionary theory, the strong divergent selection imposed by spatial variation in drug delivery leads to the emergence of a genetic polymorphism between resistant and sensitive strains supported by negative frequency dependent selection. In contrast with classic models of selection in spatially variable environments, however, this polymorphism is quasistable in the long-term, being readily undermined by compensatory evolution that tends to generate generalist resistant strains with high fitness in the absence of drugs. In other words, even spatial variation in drug selection cannot prevent resistant strains from eventually replacing sensitive ones due to the evolution of drug resistant strains with reduced costs of resistance. The implication here should be clear: in the absence of a continual pipeline of new drugs for treating infectious disease, the best we can do is slow the rate at which existing drugs lose efficacy. The use of drug sanctuaries in space may help, for a time, but natural selection will inevitably find a way to undermine our best strategies for preventing resistance.

## METHODS

### Experimental evolution

A single colony of *P. aeruginosa* strain PA14 was grown overnight in Luria-Bertrani broth (LB: bacto-tryptone 10g/L, NaCl 10 g/L, yeast extract 5 g/L) and used to found 48 populations by adding 20uL into 1.5 mL of fresh media (described below). An aliquot of the progenitor was frozen in glycerol at -80°C. Every 24 hours, a 20uL aliquot of overnight culture was added to 1.5 mL of fresh medium. Populations were propagated in 24-well microtiter plates and agitated using an orbital shaker (150 rpm) at 37°C. Samples were frozen at -80°C in glycerol every four and ten transfers, or approximately every 26 and 66 generations, respectively.

The experiment consisted of 12 replicate populations for each of four treatments growing in LB broth supplemented or not with 0.3ug/ml of the fluoroquinolone antibiotic, ciprofloxacin, for 40 days, or approximately 265 generations. This concentration of ciprofloxacin inhibits growth of the sensitive PA14 ancestor to approximately 20% of full growth in LB over a 24-hour period. Treatments were: A) a permissive environment involving daily propagation in drug-free LB (PERM); B) a constant environment with ciprofloxacin added daily (CONS); C) a temporally varying environment consisting of daily alternation in drug-free and drug-supplemented media (TEMP); and D) a spatial treatment comprised of two subpopulations, one containing drug and the other drug-free, connected through dispersal (SPAT). Dispersal was imposed by mixing 0.75mL from each pair of wells prior to transfer and inoculating a fresh pair of wells with 20uL of the combined mixture (see Supplementary Fig. S1). Samples of the mixed population were frozen. Two populations from the SPAT treatment were excluded from final analyses due to contamination.

### Minimum inhibitory concentration (MIC) and growth rate

We randomly selected eight colonies from each population at day 20, plus an additional eight colonies from the TEMP and SPAT treatments at day 40, by plating a sample of each population on agar and choosing those closest to an arbitrary point in the middle of the plate. For each isolate, we assayed resistance as the MIC to ciprofloxacin by first reviving frozen cultures overnight in LB media and then inoculating 100ul of dense culture into 96-well plates containing LB supplemented with 0.0, 0.10, 0.20, 0.39, 0.78, 1.56, 3.12, 6.25, 12.5, 25, 50, or 100ug/mL ciprofloxacin, respectively, and incubated on an orbital shaker at 37°C. Growth was scored by reading optical density (OD) at 600 nm after 48 hours. Log-transformed MICs were used for all analyses. Resistance is defined as an MIC exceeding 2ug/mL, equivalent to 4x the ancestral MIC (0.5ug/mL).

The growth of evolved SPAT isolates relative to that of PA14 was measured in LB with and without ciprofloxacin over 24 hours. In LB, 5uL of overnight culture was added to 195uL of LB and OD600 measured every 90 minutes. The same procedure was performed for ciprofloxacin growth assays, except we used 20uL of overnight culture with 180uL of LB-ciprofloxacin media (0.3ug/mL). Gen5 software (BioTek Instruments Inc., Winooski, VT) was used to estimate maximum growth rate during exponential phase for three replicates per isolate. Growth rates are expressed relative to the mean growth rate of PA14 in LB when grown on the same 96-well assay plate.

### Negative frequency dependent selection

Using reciprocal competitive invasion experiments, we estimated the strength of negative frequency dependent selection between resistant and sensitive isolates from all SPAT treatment populations that showed evidence of coexistence from the MIC assays. We first chose four random pairs of resistant and sensitive colonies from each population, then competed each pair by mixing pure culture samples from overnight cultures at ratios of 1:9, 1:1, and 9:1 by volume and allowing them to compete for two transfers (48 hours) under the same SPAT treatment transfer protocol as in the original selection experiment. We tracked the change in relative abundance of sensitive and resistant colonies by streaking 40 randomly selected colonies on 2ug/mL ciprofloxacin agar for both the initial (0 hrs) and final (48 hrs) populations. Relative fitness, w, was estimated using the equation:

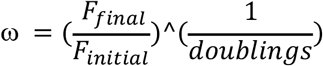

where *F*_*initial*_and *F*_*final*_ are the ratios of the frequency of resistant types to the frequency of sensitive types in the population, before and after competition, respectively. Doublings refers to the number of generations occurring between the initial and final measurements (∼13 generations).

### Dynamics of resistance

We assayed the frequency of resistance every four days for each replicate of the SPAT treatment by isolating 50 colonies at random on LB plates, streaking each colony on 2ug/mL ciprofloxacin agar, and visually inspecting each plate for growth after 24 hours at 37°C.

### Dynamics of invasion from rare

We chose a pair of resistant-sensitive isolates from population S4 at day 20 and two pairs from population S2 at day 40 to investigate the dynamics of resistance in the absence of clonal interference. These pairs were chosen because preliminary experiments indicated each had a different predicted equilibrium frequency (i.e., the frequency of resistance where ω = 1 in reciprocal competitive invasion assays). Pairs of isolates were grown overnight as pure cultures and initial ratios of resistant and sensitive strains were constructed at 1:9, 2:8, 4:6, 6:4, 8:2, and 9:1 by volume, then mixtures were propagated identically to that in the original SPAT treatment for 10 days (∼65 generations). Founding populations (day 0) and evolved populations were archived daily. The frequency of resistant individuals over time was estimated by testing the presence or absence of growth of 50 randomly selected colonies streaked on 2ug/mL ciprofloxacin agar, as described above, at days 0, 2, 4, 6, 8 and 10.

### Statistical Analysis

All statistical analyses were conducted using R Studio (Version 0.99.903; https://www.rstudio.com). We used mixed linear analyses of covariance to model the trade-off between maximum growth rate in drug free media as a function of the fixed effect of log(MIC) and day, with population treated as a random effect:
*maxV* = *log*(*MIC*)\**day* + (*log*(*MIC*)| *population*). A similar approach was used to estimate the frequency dependent fitness functions, with relative fitness (३) modeled as a function of the fixed effects of day and initial frequency of resistance (*F*_*initia*_) with random effects of isolate pair nested in population: ω = *F*_*initial*_*day + (*F*_*initiai*_ | (*population: pair*). Evidence that coexistence is supported by negative frequency dependence exists if there is both a) statistically significant negative slope between fitness and frequency of resistance, and b) the regression line crosses the relative fitness axis (i.e., ω = 1) between at a frequency between 0 and 1. Note that the *x*-intercept additionally provides an estimate of the location of the internal equilibrium frequency of resistance.

## ACKNOWLEDGEMENTS

We thank Dr. Julian Evans for help creating Figures 3A, 3B and Fig. 4. This work was funded by a Natural Sciences and Engineering Research Council (NSERC) Discovery Grant to RK and NSERC-CGSM and Ontario Graduate Scholarships to AML.

## AUTHOR CONTRIBUTIONS

Experimental design, data analysis and writing was completed by R.K. and A.M.L. Experiments were performed by A.M.L.

## SUPPLEMENTARY

**Table S1.**
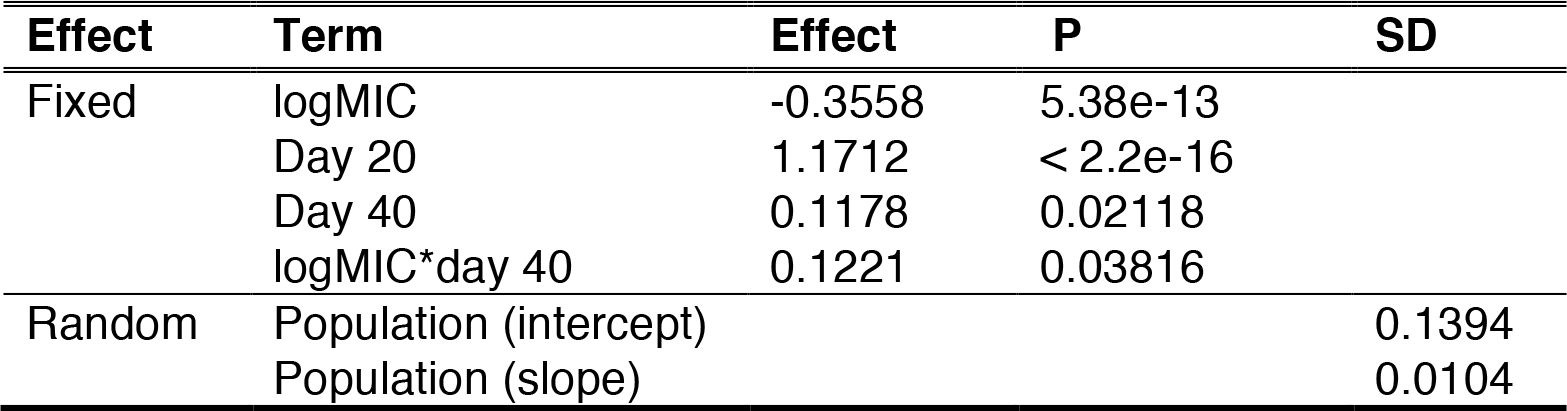
Mixed linear analysis of covariance for maximum growth rate in LB. Only populations with both resistant and sensitive isolated colonies were included for analysis.

**Table S2.** Mixed linear analysis of covariance for relative fitness (ω) of resistant types, with random effect of isolate pair nested in population. Only populations with both resistant and sensitive isolated colonies were included for analysis.

| Effect | Term | Effect | P | SD |
| --- | --- | --- | --- | --- |
| Fixed | initial frequency | -0.41871 | < 2.2e-16 |  |
|  | Day 20 | 1.124858 | < 2.2e-16 |  |
|  | Day 40 | 0.17295 | 0.005717 |  |
|  | initial frequency*day 40 | -0.06562 | 0.151243 |  |
| Random | pair : population (intercept) |  |  | 0.06733 |
|  | pair : population (slope) |  |  | 0.04819 |
|  | Population (intercept) |  |  | 0.09806 |
|  | Population (slope) |  |  | 0.04550 |

**Table S3.** Mixed linear analysis of covariance for maximum growth rate in [0.3ug/mL] ciprofloxacin. Only populations with both resistant and sensitive isolated colonies were included for analysis.

| Effect | Term | Effect | P | SD |
| --- | --- | --- | --- | --- |
| Fixed | logMIC | 0.48637 | 2.981e-09 |  |
|  | Day 20 | 0.40770 | 1.987e-12 |  |
|  | Day 40 | 0.04540 | 0.3357 |  |
|  | logMIC*day 40 | -0.03107 | 0.6167 |  |
| Random | Population (intercept) |  |  | 0.1262 |
|  | Population (slope) |  |  | 0.1842 |

**Figure S1.**
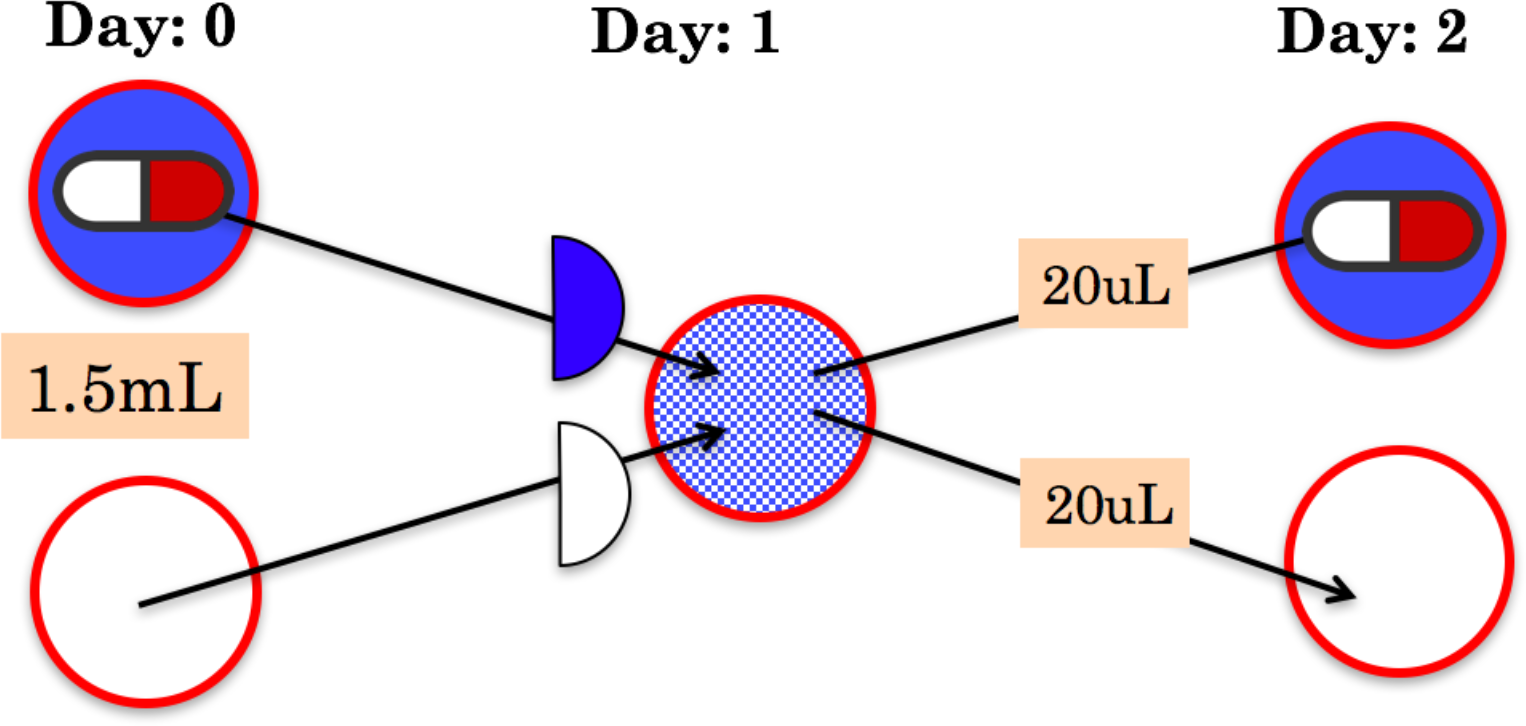
SPAT selection regime. Each population consisted of bacterial cultures propagated in two separate wells of 1.5mL LB broth and 1.5 mL ciprofloxacin(0.3ug/mL), where 0.75mL of each well was combined daily and 20uL of the combined mixture (i.e., population) was then transferred again to separate wells.

**Figure S2.**
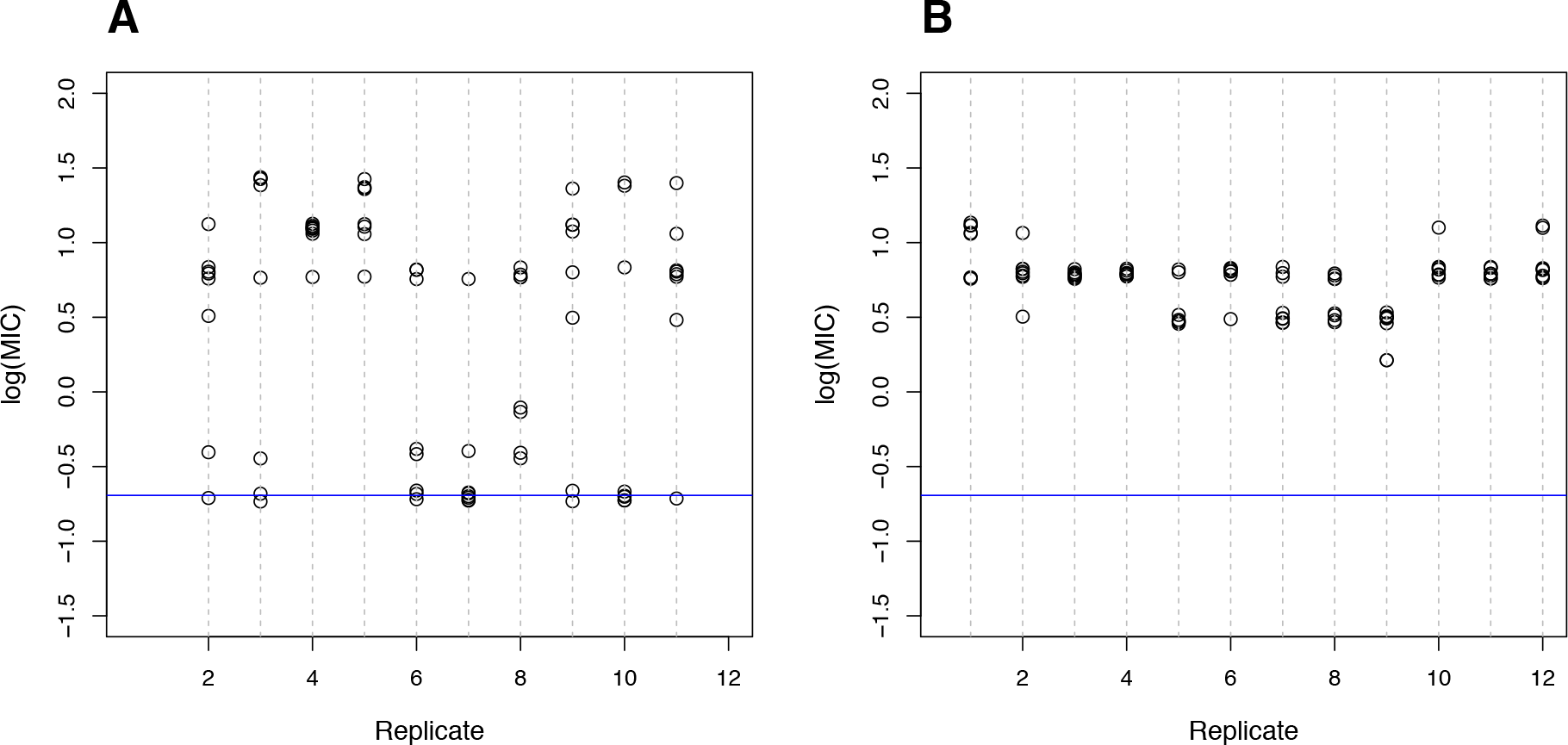
Coexistence of susceptible and resistant types maintained in SPAT treatment. Evolved levels of resistance at day 40, measured as the log minimal inhibitory concentration (MIC) of ciprofloxacin, for isolated colonies for each SPAT population **(A)** and TEMP population **(B)**. Blue axis indicates ancestral PA14 isolate resistance (MIC = 0.5ug/ml). Resistant isolates were defined as exhibiting an MIC exceeding 2ug/mL (log(MIC) = 0.3). Data points are vertically jittered for clarity.

**Figure S3.**
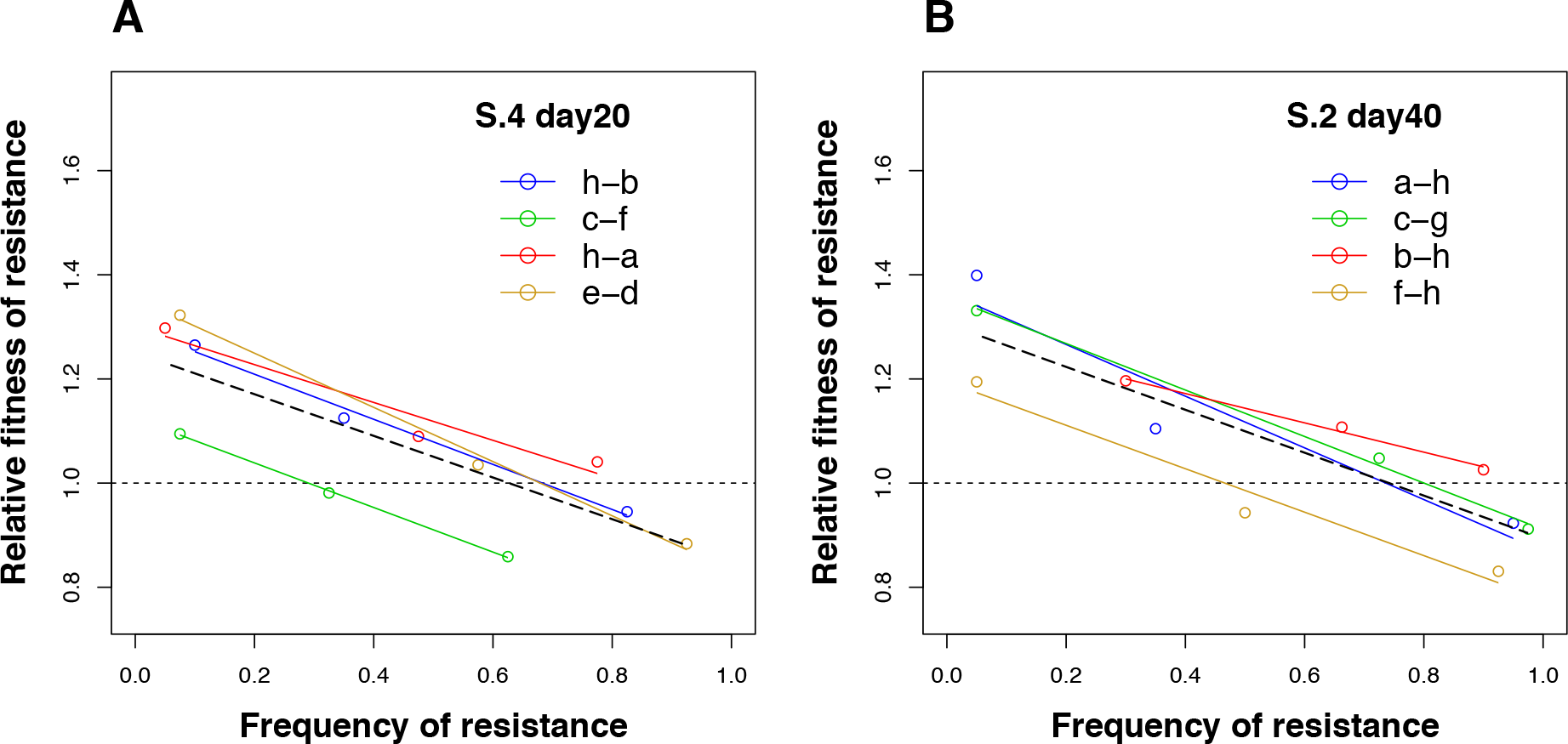
Negative frequency-dependent selection for select pairs of resistant and sensitive isolates. Fitness of a resistant colony relative to its paired sensitive colony (ω) is plotted as a function of its frequency for isolates from SPAT population S4 (day 20) **(A)**, and population S2 (day 40) **(B)**. Solid colour lines depict linear regressions of individual resistant-sensitive isolate pairs, while pooled data regressions are shown by the dashed line. The frequency of resistance at equilibrium is given by the intersection of the regression line with relative fitness (ω) = 1.

